# Geomagnetic field absence reduces adult body weight of a migratory insect by disrupting feeding behavior and appetite regulation

**DOI:** 10.1101/737361

**Authors:** Guijun Wan, Shoulin Jiang, Ming Zhang, Jingyu Zhao, Yingchao Zhang, Weidong Pan, Gregory A. Sword, Fajun Chen

## Abstract

The geomagnetic field (GMF) is well documented for its essential role as a cue used in animal orientation or navigation. Recent evidence indicates that the absence of GMF can trigger stress-like responses such as reduced body weight, as we have previously shown in newly emerged adults of the brown planthopper, *Nilaparvata lugens*. To test the hypothesis that reduced feeding in the absence of the GMF leads to a decrease of *N. lugens* body weight, we compared magnetic responses in feeding behavior, glucose levels, and expression of magnetoreception- and appetite-related genes in brown planthopper nymphs exposed to either a near-zero magnetic field (NZMF, i.e., GMF absence) or typical GMF conditions. In addition to observing the expected responses in the expression of the potential magnetosensor *cryptochromes*, the food intake of 5^th^ instar nymphs was significantly reduced in insects reared in the absence of GMF. Insects that exhibited reduced feeding reared in the absence of the GMF also had higher glucose levels which is associated with food aversion. Expression patterns of appetite-related neuropeptide genes were also altered in the absence of GMF in a manner consistent with diminishing appetite. These findings support the hypothesis that strong changes in GMF intensity can affect insect feeding behavior and underlying regulatory processes. Our results provide further evidence that magnetoreception and regulatory responses to GMF changes can affect a wide variety of biological processes.

## 1. Introduction

The geomagnetic field (GMF) is an essential cue for orientation or navigation of many migratory animals. However, the details of the biophysical mechanisms by which animals detect and respond to the GMF are poorly understood (Mouritsen, 2018). The three prevailing mechanisms of magnetoreception suggested to date for terrestrial animals are radical-pair-based magnetoreception, magnetite-based magnetoreception (Hore and Mouritsen, 2016), and electromagnetic induction detected through potential electroreceptors (Nordmann et al., 2017). Among them, cryptochrome (Cry)-mediated radical-pair-based magnetoreception has attracted the most attention, not only because it is theoretically feasible, but also because of increasing empirical support for its activity in both plants and animals (Gegear et al., 2010; Hore and Mouritsen, 2016; Kerpal et al., 2019; Maeda et al., 2012; Maffei, 2014; Ritz et al., 2000). Alternatively, theoretical hypotheses for nonspecific magnetic responses triggered by different field intensities have been proposed including the level mixing mechanism (Binhi and Prato, 2018) and dipolarly coupled three-spin mechanism (Keens et al., 2018), but currently lack any empirical support.

In a narrow sense, magnetoreception refers to the acute ability of animals to detect the GMF (Lohmann, 2010). However, increasing evidence for chronic responses of organisms to magnetic field variation suggests the importance of general magnetoreception in biological processes across taxa (Binhi and Prato, 2017; Henshaw et al., 2009; Maffei, 2014; Wan et al., 2016; Wan et al., 2014; Wan et al., 2015b; Wang et al., 2008). Among studies that subject organisms to different magnetic field intensity treatments, many concern the bioeffects induced by the near-zero magnetic field (NZMF) (Binhi and Prato, 2017; Wang et al., 2008). Although there may be the need for human beings to understand how organisms from Earth might perform during space travel where the field is ∼10,000 times weaker than that of Earth’s (Mallis and DeRoshia, 2005), the NZMF can also be taken as a sham control for the study of specific magnetoreception mechanisms (Fedele et al., 2014; Heyers et al., 2010). Indeed, organismal studies that impose a NZMF treatment do suggest the importance of GMF in maintaining organism homeostasis (Fedele et al., 2014; Van Huizen et al., 2019; Wan et al., 2014).

Studies of three closely related species of migratory rice planthoppers, *Nilaparvata lugens, Laodelphax striatellus* and *Sogatella furcifera*, have shown that the absence of the GMF (i.e., the NZMF) affects a number of physiological and behavioral processes such as nymphal development, adult longevity, body weight, wing dimorphism, positive phototaxis, and flight behaviors, potentially through Cry-mediated hormone signaling (Wan et al., 2016; Wan et al., 2014; Wan et al., 2015b). These studies highlight the GMF’s function in animal physiological homeostasis. Body weight is closely related to energy homeostasis in animals, and migratory fuelling which is characterized by gaining body mass and increased food intake during migration has been shown to be affected by GMF information (intensity, direction, or both) (Fransson et al., 2001; Henshaw et al., 2009; Henshaw et al., 2008; Kullberg et al., 2007). Interestingly, recent studies showed field intensity-dependent changes in the production of reactive oxygen species (ROS) which are regarded as signaling molecules for appetite-related neurons (Drougard et al., 2015; Sherrard et al., 2018; Van Huizen et al., 2019). Moreover, magnetic responses of adipokinetic hormone (AKH)/ AKH receptor (AKHR) signaling, reported to be involved in appetite regulation (Gáliková et al., 2017; Lin et al., 2019), was also found in adult rice planthoppers (Wan et al., 2016). Therefore, we speculated that the GMF absence-induced body weight decrease observed in rice planthoppers could be due to appetite loss and reduced feeding.

Both nanoscale magnetite particles (Pan et al., 2016) and the potential cryptochrome magnetoreceptors, Crys (Cry1 and Cry2) (Xu et al., 2016), have been systematically characterized in the migratory brown planthopper, *N. lugens*. Thus, *N. lugens* is a promising insect model for the in-depth exploration of magnetoreception mechanisms. In addition to the potential magnetoreceptor *Crys* and multifunctional *AKH*-*AKHR* signaling, here we also explored the expression of orexigenic *neuropeptide F* (*NPF*) and *short NPF* (*sNPF*) genes thought to play a role in appetite regulation (Fadda et al., 2019; Lin et al., 2019). Combining the investigation of magnetic field effects on feeding behavior along with glucose levels suggested to be negatively correlated with appetite (Timper and Bruning, 2017; Ugrankar et al., 2018), and the expression of appetite-related genes, we provide the first evidence that reducing GMF intensity negatively affects energy homeostasis in animals by altering feeding behavior and its underlying regulation.

## 2. Material and methods

### 2.1. Insect stock

The migratory adults of brown planthoppers, *N. lugens*, were collected from the paddy fields of Jiangsu Academy of Agricultural Science at Nanjing, Jiangsu province of China. The planthopper stocks were transferred to the test room and maintained on Taichung Native 1 (TN1) rice seedlings (15-30 days after planting) to expand the colony at temperature of 25±1□, 70-80% RH and 14:10 h light: dark cycle (Dark during 1800-0800 hours; All following assays were under the same environmental conditions). The Kimura B nutrient solution was used to provide nutrition for rice seedlings. The newly emerged macropterous adults were selected from the same generation of this expanded colony for the following experiments.

### 2.2. Magnetic field setups and insect exposures

Two common methods are usually utilized to mimic the absence of the GMF, either by shielding with alloys like mu-metal or through compensation with Helmholz coils. Here, we used two DC-type Helmholtz coils system (External diameter: 1200 mm) to simulate the NZMF (i.e., the GMF absence; mimic intensity: 484 ± 29 nT) and the local GMF intensity (32° 3’ 42” N, 118° 46’ 40” E; mimic intensity: 50000 ± 254 nT) at same declination (−5.2 ± 1.76°) and inclination (50.2 ± 1.19°) within the effective homogeneous areas of 300 × 300 × 300 mm^3^. The simulated magnetic fields were measured and adjusted with a fluxgate magnetometer (Model 191A, HONOR TOP Magnetoelectric Technology Co., Ltd., Qingdao, China) daily. To secure uniform environmental factors such as temperature, background disturbances, and other vibrations, the two coils systems were located in the same room with a distance of 6.5 m between each other (to avoid interference with each other). Following the previously used workflow and standards for one generation of exposure of rice planthoppers (Wan et al., 2015b). We exposed *N. lugens* individuals to NZMF vs. GMF conditions for either one generation (from mated F0 females to newly emerged F1 adults) or less than one generation (from mated F0 females to older nymphs) because their long-distance migration is usually initiated after adults emerge following at least one generation of local reproduction (Cheng et al., 2003).

### 2.3. Developmental periods of nymphs and body weight of newly emerged adults

Instar durations to the adult stage (instars 1-5) were measured in days as previously described for the white-backed planthopper, *Sogatella furcifera* (Wan et al., 2015b). Differentiating between the sexes of juveniles is very difficult and was not attempted for the molecular, behavioral, and biochemical assays of nymphs described below. Once the F1 adults emerged, individual *N. lugens* from the NZMF and GMF groups were identified to sex and weighed using Mettler Toledo XP2U precision scales (Mettler Toledo International Inc., Columbus, OH, U.S.A.) with an accuracy of ±0.1 μg.

### 2.4. Electrical penetration graph (EPG) assay for feeding behavior of nymphs

The EPG assay was conducted to monitor the feeding behavior of the 5th instar nymphs. The test arena was set up inside a Faraday cage using a GIGA-8 DC EPG amplifier system (EPG system, Wageningen University) within the effective homogeneous area of the simulated magnetic fields. In case of magnetic field anomaly, an aluminum holder was used to supply the test channel input terminals, and nymph individuals were wired with gold wires and conductive silver glue (diameter: length□=□18.5□μm: 3□cm) on their dorsum. Fifth instar nymphs randomly sampled at different times after molting were used in the EPG assays. Nymphs were starved for one hour before the test, and EPG was continuously recorded for six□hours after placing the insect on a plant for feeding. The start time for the EPG assay was the same as the daily sampling time for gene expression analysis (starved from 0900-1000 hours, and tested from 1000-1600 hours which is in the middle of the light period). Fifteen replicate EPG trials under the NMZF vs. GMF were conducted. Four□continuous hours of EPG data starting from the beginning of feeding were analyzed with EPG Stylet□+□a software (Wageningen Agricultural University, 2012). Other details of the EPG assay follow our previous work with rice planthoppers (Wan et al., 2015a).

### 2.5. Measurement of glucose content in nymphs

The hemolymph glucose content of the 5th instar nymphs was determined using commercial assay kits (Glucose assay kit; Jiancheng Bioengineering Institute, Nanjing, China) according to the manual. Nymphs were separately sampled 24h and 48h post-molt. Insects were starved for one hour before the test, and fifteen individual nymphs were randomly pooled as one sample for each sampling time with six repeats (frozen at the same time of 1000 hours by liquid nitrogen).

### 2.6. Transcript expression analysis of magnetoreception-related and orexigenic neuropeptide genes

The potential magnetoreceptor *Crys* (*Cry1* and *Cry2*), and genes mediating Adipokinetic hormone (*AKH, AKHR*) and neuropeptide Y (NPY)-like (*NPF, sNPF*) signaling which is involved in appetite regulation were selected for transcript expression analysis by quantitative real-time polymerase chain reaction (qRT-PCR) assay. Insects were separately tested as both 4th and 5th instar nymphs, with samples collected 24h and 48h post-molt for both instars. Insects were starved for one hour and 10 individuals were randomly pooled as one sample for each sampling time with six replicates. (frozen at the same time of 1000 hours by liquid nitrogen). RNA isolation, reverse transcription, and qRT-PCR assay were conducted following previous work (Wan et al., 2016). The reference genes in the transcript expression analyses were *RPL5* and *18S*, and all primer information for this study are listed in Table S1.

### 2.7. Statistical analysis

All data were analyzed using SPSS 20 (IBM Inc., Armonk, NY, U.S.A.). In this study, all outcomes were directly separated by sampling time (developmental stage) or sex to investigate the main effect of the magnetic field. Therefore, we didn’t use sampling time (developmental stage) or sex as the fixed factor in general linear models, and no post-hoc multiple comparison tests for sampling time (developmental stage) were performed. Before the analysis, data were tested for normality and homoscedasticity of variances using the Shapiro-Wilk test (*P* > 0.05) and Levene’s test (*P* > 0.05), respectively. If those conditions were satisfied, a parametric one-way analysis of variance (ANOVA; *P* < 0.05) was employed, if not, a two-tailed nonparametric Mann-Whitney *U* test was applied to determine differences in measured variables between experimental groups for female, male, each sampling time (developmental stage) or each specific feeding waveform. Effect sizes were estimated using partial *η*^2^ and Cohen’s *d* for ANOVA (small effect: partial *η*^2^ = 0.01; medium effect: partial *η*^2^ = 0.06; large effect: partial *η*^2^ = 0.14) and Mann-Whitney *U*-test (small effect: *d* = 0.2; medium effect: *d* = 0.5; large effect: *d* = 0.8), respectively, based on the benchmarks of Cohen (Cohen, 2013).

## 3. Results and discussions

### 3.1 Nymphal duration and newly-emerged adult body weight

Consistent with our previous report [17], GMF absence one again affected nymphal development and body weight of newly emerged adult *N. lugens*. In the current study we found that GMF absence significantly prolonged duration of the 4th (+34.19%; *U*_*1, 110*_ = 687.00, *P* < 0.001, *d* = 1.10) and 5th (+16.96%; *U*_*1, 110*_ = 1124.50, *P* = 0.008, *d* = 0.50) instars. It also significantly decreased body weight of newly emerged female (−14.67%; *F*_1, 47_ = 24.23, *P* < 0.001, partial *η*^2^ = 0.34) and male (−13.17%; *F*_1, 68_ = 24.14, *P* < 0.001, partial *η*^2^ = 0.26) adults (Table 1). These results suggested that the GMF is important for the maintainance of normal physiological homeostasis, and that the 4th and 5th instar nymphs were the most strongly-affected juvenile stages. Therefore, we proceeded to explore how changes in 4th and 5th instar juvenile feeding behavior and physiology were affected by dramatic changes in GMF intensity leading to the observed changes in adult weight.

**Table 1.** Nymphal duration and the newly emerged adult body weight of *Nilaparvata lugens*, under the near-zero magnetic field (NZMF, i.e., GMF absence) vs. geomagnetic field (GMF).

| Parameters | GMF | NZMF |
| --- | --- | --- |
| Nymphal duration (days)<sup>*</sup> |  |  |
| 1st instar | 3.28 ± 0.11 a | 3.17 ± 0.13 a |
| 2nd instar | 2.87 ± 0.09 a | 2.71 ± 0.10 a |
| 3rd instar | 2.98 ± 0.11 a | 2.98 ± 0.09 a |
| 4th instar | 2.72 ± 0.07 b | 3.65 ± 0.14 a |
| 5th instar | 3.43 ± 0.14 b | 4.00 ± 0.12 a |
| Adult body weight (mg)<sup>**</sup> |  |  |
| Female | 2.025 ± 0.041 a | 1.728 ± 0.044 b |
| Male | 1.595 ± 0.029 a | 1.385 ± 0.031 b |

<sup>\*</sup> Nymphal duration: n = 52 and 60 for each nymphal stage under NZMF and GMF, respectively.
<sup>\*\*</sup> Adult body weight: n = 25 and 33 for female and male under NZMF, respectively. n = 24 and 37 for female and male under GMF, respectively. Newly emerged adults were measured for adult body weight.
Note: Different lowercase letters show significant differences between the GMF and NZMF tested respectively by two-tailed Mann-Whitney *U*-test for the nymphal duration at the same developmental stage, and by one-way ANOVA for the female or male adult body weight at $P < 0.05$ .

### 3.2 EPG analysis of 5th instar feeding behavior

Given that reduced feeding in the absence of the GMF as nymphs could provide a straightforward explanation for the observed reduction in body weight of newly emerged adult *N. lugens*, we measured the feeding behavior of the 5th instar nymphs. As expected, a significant decrease in the N4b feeding waveform duration (−32.02%; *F*_*1, 28*_ = 4.93, *P* = 0.035, partial *η*^2^ = 0.15), which indicates decreased phloem ingestion (Seo et al., 2009), reflects reduced feeding in the 5th instar nymphs under the NZMF vs. GMF. In addition to the main feeding intake waveform N4b, both the duration (+48.54%; *F*_*1, 28*_ = 4.37, *P* = 0.046, partial *η*^2^ = 0.14) and frequency (−43.41%; *U*_*1, 28*_ = 170.00, *P* = 0.016, *d* = 0.97) of feeding pathway phase P as well as the frequency of xylem feeding waveform N5 (−66.67%; *U*_*1, 28*_ = 164.00, *P* = 0.033, *d* = 0.85; Figure 1) were also significantly altered in the GMF absence vs. GMF (Seo et al., 2009). These responses may be due to potential magnetic field effects on salivary sheath-involved regulators (Huang et al., 2015), but we did not directly test for this possibility.

**Figure 1.**
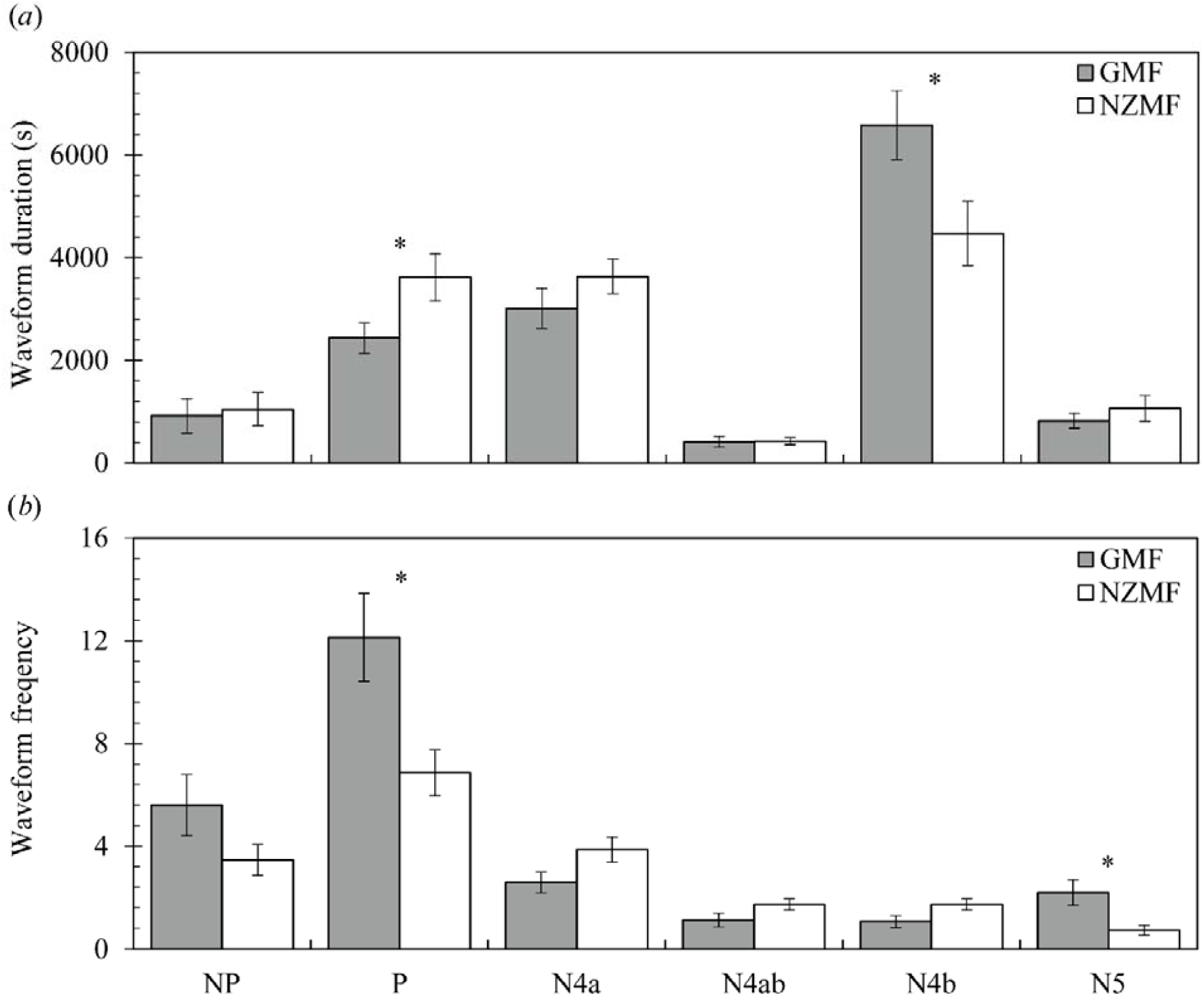
The geomagnetic field (GMF) absence altered both duration (a) and frequency (b) of specific feeding waveforms in 5th instar nymph measured through the electrical penetration graph assay. NZMF, near-zero magnetic field (i.e., the GMF absence); NP, non-penetration waveform; P, pathway phase, the sum of irregular mixed and transition phase prior to N4a; N4a, sieve element salivation waveform; N4ab, transition phase between N4a and N4b; N4b, phloem ingestion waveform; N5, xylem feeding waveform. N□=□15 for each treatment and 4 hours of feeding behavior were analyzed for each nymph. Only significant differences between the GMF and NZMF are marked with asterisk tested by one-way ANOVA for the duration or by two-tailed Mann-Whitney *U*-test for the frequency of specific feeding waves at *P*□<□0.05.

### 3.3 Glucose levels of 5th instar nymphs

*Drosophila* larvae with increased glucose levels have an aversion to feeding, and glucose has been proposed as a potential negative regulator of appetite (Ugrankar et al., 2018). In this study, elevated glucose levels were found in 5th instar nymphs at both 24h (+16.98%; *F*_*1, 10*_ = 9.54, *P* = 0.011, partial *η*^2^ = 0.49) and 48h (+20.05%; *F*_*1, 10*_ = 9.86, *P* = 0.011, partial *η*^2^ = 0.50; Figure 2) after molting under the NZMF vs. GMF. This finding is consistent with the decreased phloem ingestion by the 5th instar nymphs in response to the GMF absence (i.e., the NZMF) that we also observed. Combining the above behavioral and physiological results, we further speculated that the normal regulatory processes underlying feeding behavior of *N. lugens*, such as appetite regulation, might be disrupted by the absence of the GMF. Hence, temporal transcript expression of genes related to appetite regulation were investigated in both 4th and 5th instar nymphs.

**Figure 2.**
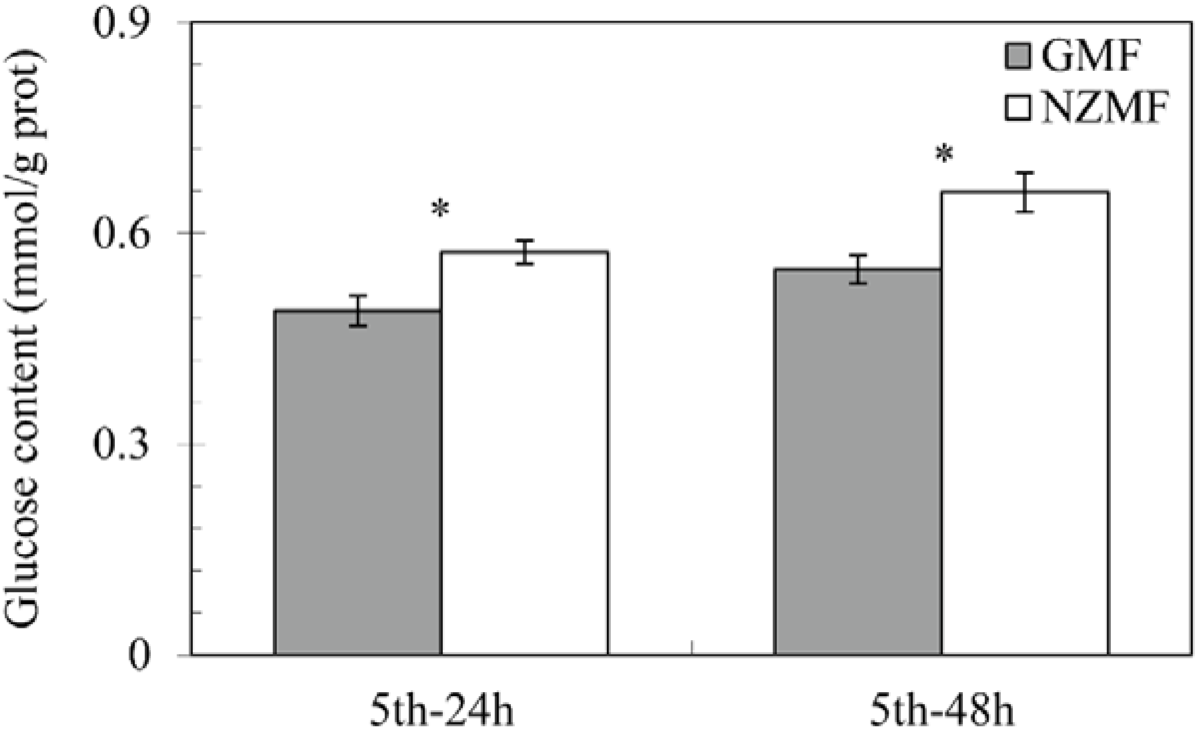
The geomagnetic field (GMF) absence increased the glucose content of 5th instar nymphs. NZMF, near-zero magnetic field; mmol/g prot, mmol/g protein. Fifteen individual nymphs were randomly mixed as one sample for each sampling time with six repeats. The columns represent averages with vertical bars indicating SE. Only significant differences between NZMF and GMF at the same sampling time are marked with asterisks tested by the one-way ANOVA at *P* < 0.05.

### 3.4 Temporal transcript expression of magnetoreception-related *Cryptochromes* and appetite-related genes at 24h and 48h after molting into the 4th or 5th instar

We first investigated the temporal transcript expression of multifunctional *Cry1* and *Cry2*. As expected, the *Drosophila-like Cry1* (Gegear et al., 2010; Kutta et al., 2017), which is the posited magnetosensor, showed significantly up-regulated gene expression at 48h after molting into the 4th (+22.41%; *F*_*1, 10*_ = 11.03, *P* = 0.008, partial *η*^2^ = 0.52) and 5th (+27.12%; *F*_*1, 10*_ = 14.34, *P* = 0.004, partial *η*^2^ = 0.59) instar under the NZMF compared with the GMF. We also found significantly altered *Cry2* expression pattern under the NZMF at 24h after molting into the 4th (+32.32%; *F*_*1, 10*_ = 22.02, *P* = 0.001, partial *η*^2^ = 0.69) and 5th instar (+65.43%; *F*_*1, 10*_ = 34.93, *P* < 0.001, partial *η*^2^ = 0.78), and at 48h after molting into the 5th instar (−16.65%; *F*_*1, 10*_ = 10.34, *P* = 0.009, partial *η*^2^ = 0.51; Figure 3a). Indeed, the vertebrate-like Cry2 gene has also been shown to be related to magnetoreception in cockroaches (Bazalova et al., 2016; Gegear et al., 2010). However, since it is classified as a vestigial flavoprotein (Kutta et al., 2017), and is unlikely to be the magnetoreceptor, Cry2 may work together with another chromophore (e.g., FAD in *Drosophila-like* Cry1) in magnetoreception. Moreover, given that Cry2 is a core component of the circadian clock (Jiang et al., 2018; Zhang et al., 2017), which has been shown to be sensitive to changes in magnetic field intensity (Fedele et al., 2014; Yoshii et al., 2009), the altered gene expression pattern of *Cry2* observed in our study was more likely related to its role in circadian mechanisms.

**Figure 3.**
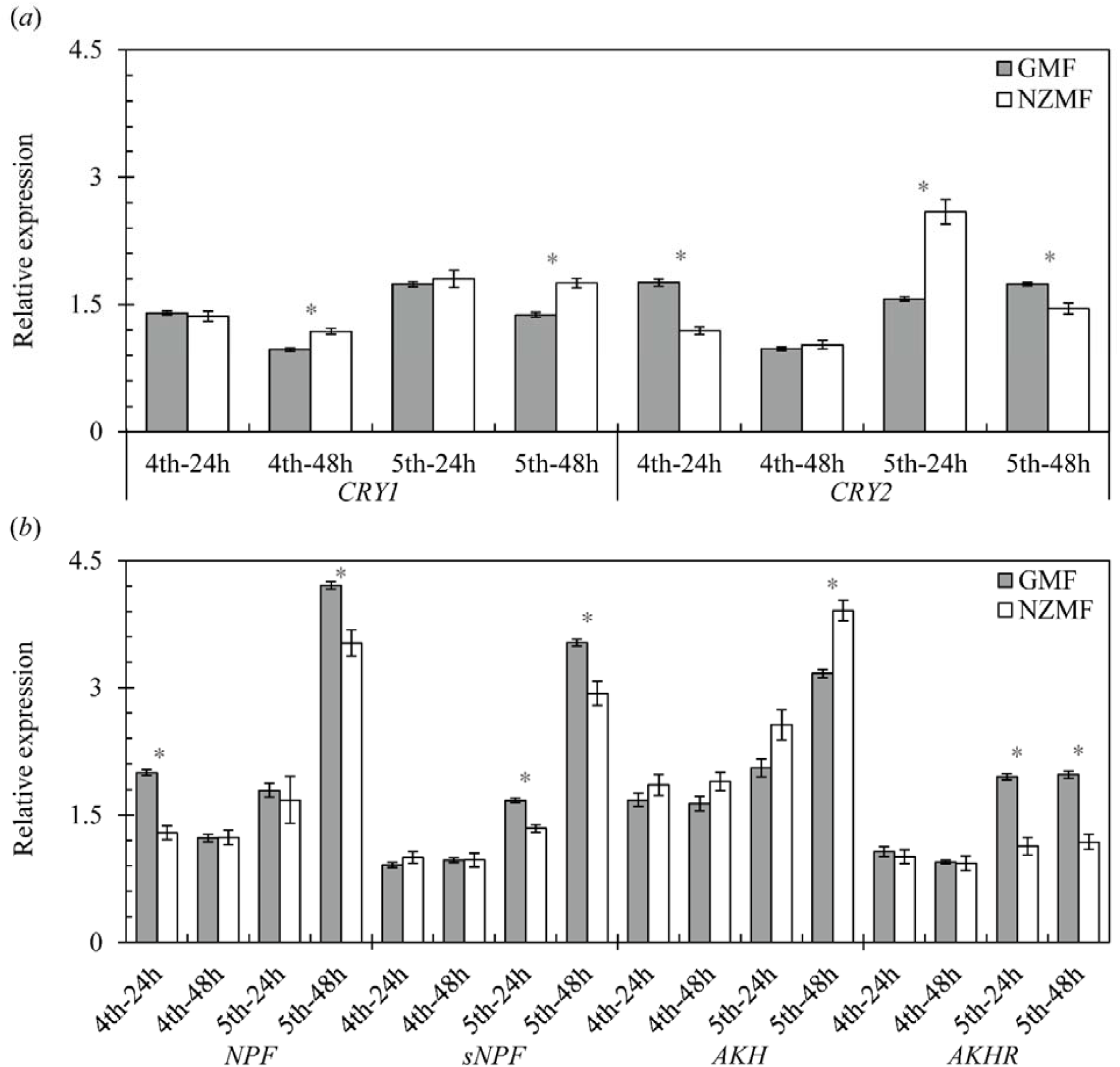
Temporal gene expression patterns of (a) magnetoreception-related *Cryptochromes* (*Cry1* and *Cry2*) and (b) appetite-related *NPF, sNPF, AKH and AKHR* at 24h and 48h after molting into the 4th and 5th instar of *Nilaparvata lugens*, under the near-zero magnetic field (NZMF, i.e., the GMF absence) vs. the geomagnetic field (GMF). Ten individual nymphs were randomly mixed as one sample for each sampling time with six repeats. The columns represent averages with vertical bars indicating SE. Only significant differences between NZMF and GMF for each gene at the same sampling time are marked with asterisks tested by the one-way ANOVA at *P* < 0.05.

Previous work in adult *S. furcifera*, a closely related planthopper species to *N. lugens*, suggested a Crys-circadian clock-AKH/AKHR signaling pathway in response to the GMF absence (Wan et al., 2016). AKH is an insect analog of the human hormone glucagon (Gáliková et al., 2017). The AKH/AKHR signaling plays crucial roles in energy mobilization (Galikova et al., 2015), antioxidative stress reactions (Kodrik et al., 2015), anti-obesity, and appetite regulation (Gáliková et al., 2017). Interestingly, our results indicated that AKH/AKHR signaling changed in response to changes in the GMF intensity in 5th instar *N. lugens* nymphs. The transcript expression of *AKH* was significantly up-regulated at 48h after molting into the 5th instar (+23.51%; *F*_*1, 10*_ = 15.28, *P* = 0.003, partial *η*^2^ = 0.60), while *AKHR*, which is a typical G protein-coupled receptor, was significantly down-regulated at both 24h (−41.93%; *F*_*1, 10*_ = 31.21, *P* < 0.001, partial *η*^2^ = 0.76) and 48h (−40.17%; *c*) after molting into the 5th instar. AKH in *Drosophila* is an orexigenic neuropeptide, and AKH deficiency by knockout of *AKH* or *AKHR* causes obesity with decreased food intake (Gáliková et al., 2017). *AKHR* silencing was also reported to cause obesity in *N. lugens*, while AKH application produced a slim phenotype (Lu et al., 2018). Therefore, in our study, significant down-regulation of *AKHR* at 24h and 48h in the 5th instar nymphs likely lead to hypophagia which is consistent with what we found in the feeding behavior assay. The *AKH* and its receptor *AKHR* may work reciprocally in the 5th instar nymphs of *N. lugens* to maintain energy homeostasis stressed by the GMF absence.

Two neuropeptide Y (NPY)-like appetite regulatory factors, *NPF* and *sNPF*, were also found sensitive to the GMF absence in both 4th and 5th instar nymphs. For *NPF*, consistent significant down-regulation of gene expression was found at 24h after molting into the 4th instar (−35.50%; *F*_*1*,_ *10* = 29.37, *P* < 0.001, partial *η*^2^ = 0.75) and at 48h after molting into the 5th instar (−16.16%; *F*_*1, 10*_ = 11.16, *P* = 0.007, partial *η*^2^ = 0.53), which is consistent with a decreased appetite in the older nymphs. Similar significant down-regulated gene expression of sNPF was found at both 24h (−19.84%; *F*_*1, 10*_ = 13.11, *P* = 0.005, partial *η*^2^ = 0.57) and 48h (−17.03%; *F*_*1, 10*_ = 10.04, *P* = 0.010, partial *η*^2^ = 0.50; Figure 3b) after molting into the 5th instar. It has been reported that sNPF, unlike NPF, acts as either a stimulating or suppressing factor in a more intricate species-specific way [25]. Given that decreased phloem ingestion of the 5th instar nymphs was found in the absence of the GMF, the sNPF in *N. lugens* should be an appetite stimulator. Taken together, magnetic field effects on glucose levels and transcript expression patterns of orexigenic neuropeptide genes in older nymphs all point to a loss of appetite triggered by the absence of the GMF, which is accordance with our expectations. Therefore, disrupted feeding behavior mediated by appetite loss in *N. lugans* nymphs under the NZMF vs. GMF could be the main reason for the decreased body weight observed in the newly emerged adults.

Although environmental factors like temperature and photoperiod have been reported to affect animal appetite (Biswas et al., 2005; Zheng et al., 2019), it might seem surprising that absence of the GMF could also disrupt appetite regulation leading to changes in feeding behavior that contribute to the adult body weight loss. However, this may be easier to understand if we link the circadian clock, AKH/AKHR signaling, and NPF/sNPF signaling (Lee et al., 2006). The potential magnetoreceptor Crys cryptochromes themselves play multifunctional roles in both magnetoreception and circadian clock mechanisms (Chaves et al., 2011), and genetic evidence has already demonstrated the magnetosensitivity of the circadian clock to magnetic field intensity changes (Fedele et al., 2014; Yoshii et al., 2009). Potential interactions between their action on magnetoreception and the circadian clock could be the driving force for the chronic magnetic responses we found here, because both AKH/AKHR and NPF/sNPF signaling, and even feeding behavior are under the regulation of the circadian clock (Bechtold and Loudon, 2013; Chatterjee et al., 2010; Lee et al., 2006). Additionally, recent computational work suggests that [FAD^·−^–HO_2_^·^] or [FADH^·^–O_2_^·−^] can be alternative radical pairs at the origin of magnetoreception in Crys (Mondal and Huix-Rotllant, 2019), and ROS levels have also been shown to be sensitive to the changes in magnetic field intensity (Sherrard et al., 2018; Van Huizen et al., 2019). Moreover, it is well documented that ROS are critical in the regulation of energy metabolism and food intake due to their versatile roles in mediating the expression of vertebrate NPY (homolog of invertebrate NPF), glucose levels (Drougard et al., 2015) as well as the AKH/AKHR pathway (Kodrik et al., 2015). Therefore, another pathway that could explain GMF-involved appetite regulation is likely to be affected by ROS.

## 4. Conclusion

We presented the first evidence in a nocturnal insect migrant that appetite-related regulatory pathways and phenotypic outcomes can be influenced by changes in GMF intensity. The changes in feeding behavior, glucose levels, and gene expression patterns induced by strong changes in GMF intensity that we observed highlight the role of the GMF in the complicated system of maintaining animal energy homeostasis. Energy homeostasis is also critical for migratory animals that can undergo large-scale spatial displacements in a short period of time (Chapman et al., 2015). A difference in GMF intensity between the emigration and immigration areas could be important environmental cues for the regulation of biological processes during migration. (e.g., fuelling behavior mediated by the GMF in several migratory birds) (Fransson et al., 2001; Henshaw et al., 2009; Henshaw et al., 2008; Kullberg et al., 2007). Future work should determine if physiological processes related to migration are also influenced by naturally-occurring levels of GMF variation in the geophysical environment.

## Funding

This work was supported by the National Natural Science Foundation of China (31701787, 31470454, 31670855 and 51037006), the natural science foundation of Jiangsu Province Youth Fund (BK20160717), the Fundamental Research Funds for the Central Universities (KJQN201820), the Nanjing Agricultural University Start-up Fund (82162045), the Jiangsu Province Postdoctoral Science Foundation (1601196C), and the National Basic Research Program of China (973) (2010CB126200).

## Acknowledgments

We thank Fanqi Liu, Ruiying Liu and Jinglan He for their help in keeping insect stocks and pilot experiment for molecular and biochemical trials, and Likun Li for maintaining the coils system.

---

**Table S1.**
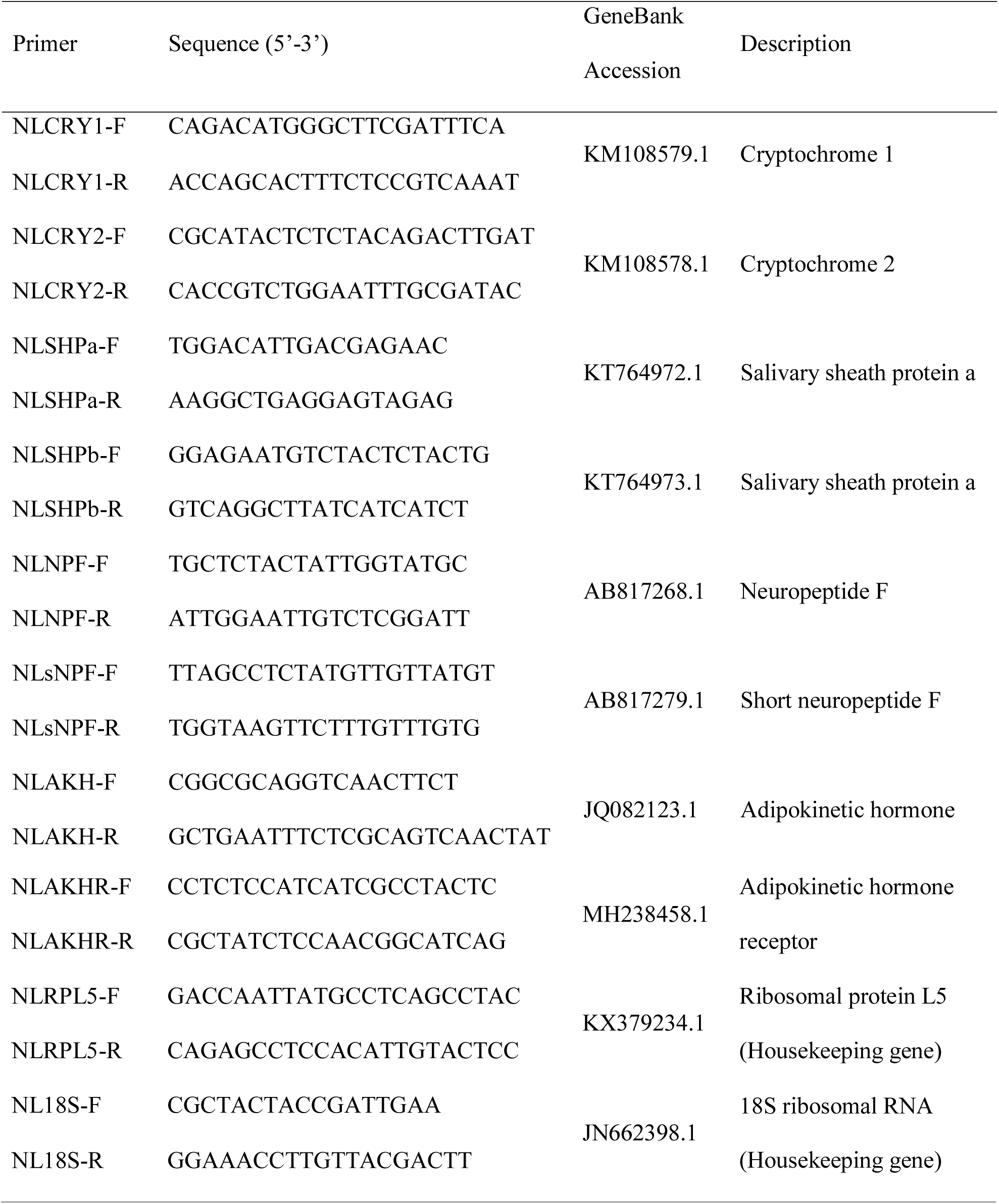
Primers used to measure the transcript expression of selected genes in the qRT-PCR experiments

