## Supplementary material for "Geomagnetic field absence reduces adult body weight of a migratory insect by disrupting feeding behavior and appetite regulation": Table S1

**Table S1** Primers used to measure the transcript expression of selected genes in the qRT-PCR experiments

| Primer | Sequence (5’-3’) | GeneBank Accession | Description |
| --- | --- | --- | --- |
| NLCRY1-F | CAGACATGGGCTTCGATTTCA | KM108579.1 | Cryptochrome 1 |
| NLCRY1-R | ACCAGCACTTTCTCCGTCAAAT |  |  |
| NLCRY2-F | CGCATACTCTCTACAGACTTGAT | KM108578.1 | Cryptochrome 2 |
| NLCRY2-R | CACCGTCTGGAATTTGCGATAC |  |  |
| NLSHPa-F | TGGACATTGACGAGAAC | KT764972.1 | Salivary sheath protein a |
| NLSHPa-R | AAGGCTGAGGAGTAGAG |  |  |
| NLSHPb-F | GGAGAATGTCTACTCTACTG | KT764973.1 | Salivary sheath protein a |
| NLSHPb-R | GTCAGGCTTATCATCATCT |  |  |
| NLNPF-F | TGCTCTACTATTGGTATGC | AB817268.1 | Neuropeptide F |
| NLNPF-R | ATTGGAATTGTCTCGGATT |  |  |
| NLsNPF-F | TTAGCCTCTATGTTGTTATGT | AB817279.1 | Short neuropeptide F |
| NLsNPF-R | TGGTAAGTTCTTTGTTTGTG |  |  |
| NLAKH-F | CGGCGCAGGTCAACTTCT | JQ082123.1 | Adipokinetic hormone |
| NLAKH-R | GCTGAATTTCTCGCAGTCAACTAT |  |  |
| NLAKHR-F | CCTCTCCATCATCGCCTACTC | MH238458.1 | Adipokinetic hormone receptor |
| NLAKHR-R | CGCTATCTCCAACGGCATCAG |  |  |
| NLRPL5-F | GACCAATTATGCCTCAGCCTAC | KX379234.1 | Ribosomal protein L5  (Housekeeping gene) |
| NLRPL5-R | CAGAGCCTCCACATTGTACTCC |  |  |
| NL18S-F | CGCTACTACCGATTGAA | JN662398.1 | 18S ribosomal RNA  (Housekeeping gene) |
| NL18S-R | GGAAACCTTGTTACGACTT |  |  |
